# Genetic and epigenetic contributions to variation in transposable element expression responses to abiotic stress in maize

**DOI:** 10.1101/2020.08.26.268102

**Authors:** Zhikai Liang, Sarah N. Anderson, Jaclyn M. Noshay, Peter A. Crisp, Tara A. Enders, Nathan M. Springer

## Abstract

- Transposable elements (TEs) pervade most eukaryotic genomes but the repetitive nature of TEs has complicated the analysis of their expression. Although the majority of TEs are silent, we document the activation of some TEs during abiotic stress.
- TE expression was monitored in seedling leaf tissue of maize inbreds subjected to heat or cold stress conditions. DNA methylation profiles and comparative genomics were used to probe the variability of TE expression responses.
- Although there was no evidence for a genome-wide activation of TEs, a subset of TE families generate transcripts only in stress conditions. There is substantial variation for which TE families exhibit stress-responsive expression in the three genotypes. The stress-responsive activation of a TE family can often be attributed to a small number of elements in the family. These elements that are activated often contain small regions lacking DNA methylation, while fully methylated elements are rarely expressed. A comparison of the expression of specific TEs in different maize genotypes reveals high levels of variability that can be attributed to both genome content differences and epigenetic variation.
- This study provides insights into the genetic and epigenetic factors that influence TE regulation in normal and stress conditions.

## Introduction

Transposable elements (TE) are the predominant portion of many plant genomes. These mobile elements can accumulate to quite high copy through a variety of mechanisms depending on the specific class or type of TE (Wicker *et al.*, 2007). Class I TEs utilize RNA intermediates for transposition while class II elements utilize DNA intermediates. A structural annotation of TEs in the maize genome reveals >150,000 class I and class II elements but the class I elements tend to be much longer and account for a substantially larger portion of the genome (Stitzer *et al.*; Jiao *et al.*, 2017). The majority of these TEs are transcriptionally silent and are found in chromatin containing high levels of DNA methylation and histone modifications associated with heterochromatin (Rabinowicz *et al.*, 1999). However, a subset of the TEs are transcriptionally active with tissue-specific expression (Anderson *et al.*, 2019b).

Transcripts arising from TEs can have diverse functional consequences. In some cases, the transcripts can encode the RNA and proteins that are necessary for TE mobilization and lead to transposition. In other cases, the transcripts can provide outward reading promoters that result in novel expression patterns or isoforms for nearby protein coding genes. Transcripts arising from TEs could also reflect cryptic promoters within the TE that respond to enhancers or other regulatory elements within or near the TE. The transcripts arising from TEs can result in sRNAs that may trigger silencing of other TEs from the same family or genes with related sequences. The diverse potential influences of TEs on the expression or regulation of genes may be co-opted to provide novel regulatory responses.

Prior research has found that some plant TEs can be transcriptionally activated by environmental stress (Wessler, 1996; Negi *et al.*, 2016; Galindo-González *et al.*, 2017; Lanciano & Mirouze, 2018; Benoit *et al*., 2019). These include examples of TEs activated during tissue culture (Peschke *et al.*, 1987; Hirochika, 1993), heat stress (Ito *et al.*, 2011; Cavrak *et al.*, 2014), cold stress (Jiang *et al.*, 2004) or biotic stresses (Grandbastien *et al.*, 2005). While many of these examples were identified through detection of novel TE insertions or targeted assays for activity, the advent of transcriptome profiling has provided opportunities to survey all TEs. However, the highly repetitive nature of most TEs leads to complications in monitoring expression of these elements. Monitoring the expression of a family can allow for multi-mapping reads but does not identify the specific elements that contribute transcripts (Anderson *et al.*, 2019b).

Prior research has identified highly dynamic expression of maize TEs in different tissues or development stages (Vicient *et al*., 2010; Anderson *et al.*, 2019b) and has suggested that some maize TE families may be associated with triggering stress-responsive expression for nearby genes (Makarevitch *et al.*, 2015). In this study, we apply approaches for monitoring both per-family and per-element expression of TEs to document the transcriptional response of maize TEs to heat or cold stress in seedling leaf tissue. We find that many TEs exhibit expression changes in response to heat or cold and that these responses are highly variable among genotypes. This variation among genotypes includes examples of varying genomic content of specific elements as well as examples of elements that are present in multiple genomes with distinct transcriptional responses.

## Materials and Methods

### Plant materials and conditions

Seeds from three maize genotypes, B73, Mo17 and W22, were imbibed overnight and then planted in soil. Seedlings were grown in the growth chamber at University of Minnesota for 13 days, with water added every 2 days. Plants were grown under 16 hours of light with a temperature of 30 °C in the day and 20 °C at night. On day 13 when the lights turned on, heat and cold stressed plants were moved to incubators while control plants remained in consistent conditions. The cold stress incubator was set to 6 °C and the heat stress incubator was set to 42 °C. All plants were removed after 4 hours, and half of the 3rd leaf was cut longitudinally from four plants was pooled for each replicate. Every genotype under each treatment contains five biological replicates, except that B73 under control condition contains four biological replicates.

### RNA-seq data processing

RNA-seq reads were generated using NovaSeq 6000 in paired-end 150 bp mode and deposited in the NCBI SRA database under the accession ID PRJNA657262. Raw reads per sample were trimmed and preprocessed using trim_galore in default settings (version 0.6.4; http://www.bioinformatics.babraham.ac.uk/projects/trim_galore/). Preprocessed reads per sample in each genotype (B73, W22 and Mo17) were aligned to their own indexed reference genome (B73 AGPv4, W22 and Mo17) using hisat2 (Kim *et al.*, 2015) with up to 20 multi-mapping positions (version 2.1.0; -k 20 --no-mixed --no-discordant) respectively. Alignment files were converted to bam format using samtools (version 1.9) (Li *et al.*, 2009).

### Gene and TE expression calling

Both genes and TE elements on chromosomes were retained for further analysis. Filtered TE annotations of B73 AGPv4, W22 and Mo17 in disjoined modes (https://github.com/SNAnderson/maizeTE_variation) were used for calculating TE expression. Gene annotation of B73 in version 4.41 was downloaded from Ensemble, and W22 and Mo17 were downloaded from the MaizeGDB database. Gene exon regions were subtracted from TE annotation due to ambiguous mapping reads in overlapped regions between genes and TEs. Gene body regions per genotype in gene annotation files were appended to subtracted TE annotation files respectively. Same annotation files were used to check intersected regions with unmethylated regions per genotype. Raw unique read counts per gene and TE element were calculated using HTSeq (Anders *et al.*, 2015) (version 0.11.2; -s no -m union -a 0). Because TE elements were clustered into family levels based on their sequence similarity within each family, multi-mapping reads assigned to each TE family were also considered and summed with unique mapping reads per TE element to give per-family TE counts (Anderson *et al.*, 2019b). In summary, three count files --genes, TE elements and TE families --were used in the analysis. Counts per million mapping reads (CPM) per gene/TE element/TE family were calculated in each sample separately. For each genotype, features (genes, TE elements and TE families) were considered as expressed in each abiotic stress experiment when mean CPM greater than 1 in either “control + cold” or “control + heat”. Differentially expressed features were calculated using the DESeq function in R package DESeq2 for cold or heat conditions. Upregulated features were defined as log2FoldChange > 1 with adjusted p-value <= 0.05. Downregulated features were defined as log2FoldChange < −1 with adjusted p-value <= 0.05. Non-upregulated features were defined as expressed features (mean CPM > 1) that were not upregulated.

Total expressed TE elements were considered as expressed TE elements in either cold or heat experiment. Cold specific TE elements were determined as mean CPM value of replicates in cold condition > 1 and mean CPM value of replicates in both control and heat condition <= 0.1. The same parameter was applied on defining heat specific TE elements.

### Sample clustering

Using inverse hyperbolic sine (asinh) transformed CPM values of expressed genes/TE families/TE elements, the principal component analysis was performed using the prcomp function in R. The first two principal components were used to separate analyzed samples.

### GO enrichment for upregulated genes

Aggregate GO terms without duplication and redundancy were downloaded from the maize GAMER project (Wimalanathan *et al.*, 2018) (https://dill-picl.org/projects/gomap/gomap-datasets/) for B73 AGPv4, W22 and Mo17 gene models. GO enrichment analysis was performed using goatools (Klopfenstein *et al.*, 2018) for upregulated genes per genotype under cold/heat stress condition and p-value per GO term was corrected using Bonferroni multiple testing correction methods. GO terms with corrected p-value less than 0.05 were reported.

### Unique mapping ratio per TE family

For either cold or heat experiment, the unique mapping ratio per TE family was calculated using the mean value of raw multiple read counts per sample divided by the mean value of raw unique read counts per TE family in the identical sample per experiment.

### Gene and TE correspondence across genotypes

Homologous genes across B73, W22 and Mo17 were generated using three complementary approaches (SynMap (Lyons *et al.*, 2008) + MUMmer (Kurtz *et al.*, 2004) + OrthoFinder (Emms & Kelly, 2015)) in an iterative manner (Anderson *et al.*, 2019a). As duplicated genes in one genome could be assigned to one gene in another genome in this list, only the one of duplicated genes with the highest LASTZ (http://www.bx.psu.edu/miller_lab/dist/README.lastz-1.02.00/README.lastz-1.02.00a.html) alignment score compared to the gene in another genome was retained to produce a 1:1 syntenic gene list between two of three maize genomes. A set of non-redundant TE elements was used to identify correspondence of individual TE elements in B73, W22 and Mo17 (Anderson *et al.*, 2019a). TE families between genomes have matched the first eight letters and then were considered as shared.

### Phylogenetic tree construction of TE elements within a TE family

For a selected LTR retrotransposon family, the 5' LTR sequence was used to represent each individual TE element. The multiple alignment file was produced using Mafft (version 7.464) L-INS-i algorithm (Katoh & Standley, 2013). FastTree (version 2.1.10) (Price *et al.*, 2010) was used to generate the maximum likelihood phylogenetic tree. The tree structure per selected TE family was visualized using the R package ggtree (Yu *et al.*, 2017).

### Identification of unmethylated regions

In this study we utilized whole-genome bisulfite sequencing of a subset of the same seedling leaf samples from 3 maize inbreds (B73, Mo17, W22) and 2 stress conditions (heat and cold) as described in plant materials and conditions. DNA extractions were performed using the DNeasy Plant Mini kit (Qiagen). 1ug of DNA in 50ug of water was sheared using an Ultrasonicator (Covaris) to approximately 200-350 bp fragments. 20ul of sheared DNA was then bisulfite converted using the EZ DNA Methylation-Lightning Kit (Zymo Research) as per the manufacturer’s instructions and eluted in a final volume of 15ul. Then 7.5ul of the fragmented, bisulfite-converted sample was used as input for library preparation using the ACCEL-NGS Methyl-Seq DNA Library Kit (SWIFT Biosciences). Bisulfite libraries were sequenced on a HiSeq 2500 in paired-end mode at the University of Minnesota Genomics Center. Output has been deposited in the NCBI SRA database under accession number PRJNA657677.

Trim_galore (Krueger, 2012) was used to trim adapter sequences and additional sequence noise, read quality was assessed with the default parameters and paired-end read mode.

Reads that passed quality control were aligned to the corresponding genome assembly. Alignments were conducted using BSMAP-2.90 (Xi & Li, 2009) allowing up to 5 mismatches and a quality threshold of 20 (-v 5 -q 20). Duplicated reads were detected and removed using picard-tools-1.102 (http://broadinstitute.github.io/picard/) and SAMtools (Li *et al.*, 2009). The resulting alignment file, merged for all samples with the same tissue, condition and genotype, was then used to determine methylation level for each cytosine using BSMAP tools. The *methylratio.py* script from *bsmap v2.74* was used to extract per site methylation data summaries for each context (CH/CHG/CHH) and reads were summarized into non-overlapping 100bp windows tiling the genome.

Unmethylated regions were determined following methods described in Crisp et al (2020). In brief, each 100bp tile was classified as missing data, RdDM, Heterochromatin, CG only, Unmethylated, or Intermediate according to their methylation state in each context. Unmethylated tiles were those which contained CG, CHG, and CHH levels less than 10%. Adjacent unmethylated tiles (UMTs) were merged and those larger than 300bp were kept and defined as unmethylated regions (UMRs).

### Overlap between unmethylated and partial expressed regions in TE elements

For each genotype, UMR regions were subtracted from regions of expressed TE element (TE expressed in either cold or heat experiment), cold specific TE elements or heat specific TE elements. For each selected UMR region, ten of 100 bins were extended from both sides and counted the number of unique mapping reads per bin. Any bin overlapped with another UMR will be labelled as missing counts. In total, each UMR would produce 21 bin regions, including 10 5’ extended bins, 1 UMR region and 10 3’ extended bins. For each UMR and its extended 20 bins, their overlapped values were normalized between 0 and 1 to draw heatmaps. Overall, mean normalized values of each bin of all UMR regions were calculated to visualize line plots.

## Results

### Many TEs are activated by abiotic stress in maize

Assessing the expression of TEs is complicated by the highly repetitive nature of the individual TEs within a family. In order to broadly survey the expression of TEs, we initially focused on monitoring the per-family expression of TEs. We implemented an approach that counts all unique and multi-mapping reads that map to members of a TE family (Anderson *et al.*, 2019b). This per-family expression estimate allows us to monitor all TE families even if they are highly repetitive which provides the opportunity to quantify the amount of TE transcripts in RNAseq datasets. However, this per-family expression approach does not specify the amount of expression from individual elements.

The per-family expression of TEs was assessed in a novel dataset that includes at least four biological replicates of seedling leaf material from plants subjected to control, heat or stress conditions and includes three genotypes (B73, Mo17 and W22) with *de novo* genome assemblies and consistent TE annotations (Stitzer *et al.*; Jiao *et al.*, 2017; Springer *et al.*, 2018; Sun *et al.*, 2018; Anderson *et al.*, 2019a). We identified genes and TE families that exhibit significant expression changes in the abiotic stress relative to control conditions for all three genotypes (Figure 1a). A GO enrichment analysis of up-regulated genes reveals enrichment for terms associating with abiotic stress responses (GO:0009409 "response to cold" in cold stress; GO:0009408 "response to heat" in heat stress), suggesting effective cold/heat responses have been triggered in these plants (Table S1). In general, the proportion of TE families that exhibit up- or down-regulation in cold stress is quite similar to the proportion of genes with differential expression (Figure 1a). In heat-stress there are more examples of up-regulation than down-regulation of TEs (Figure 1a). A PCA analysis performed for each genotype revealed that using genes or TE per-family expression levels clustered the samples based on treatment (Figure S1). The heat stress resulted in a greater difference relative to the control than the cold stress for all genotypes, and there was not a major difference in the differentiation between conditions when using genes compared to TE expression (Figure S1). The up-regulated TE families include examples from each superfamily of TEs (Table S2). Expressed TE families are enriched for families with large numbers of elements compared to the genome-wide set of TE families (Figure S2). However, the up-regulated TE families do not show strong enrichments for large or small TE families compared to other expressed TEs (Figure S2). Overall, there are a substantial number of TE families that respond to abiotic stress but the majority of TE families do not exhibit changes in expression level in these stress conditions suggesting a specific, rather than global, response of TE families to abiotic stress.

**Figure 1.**
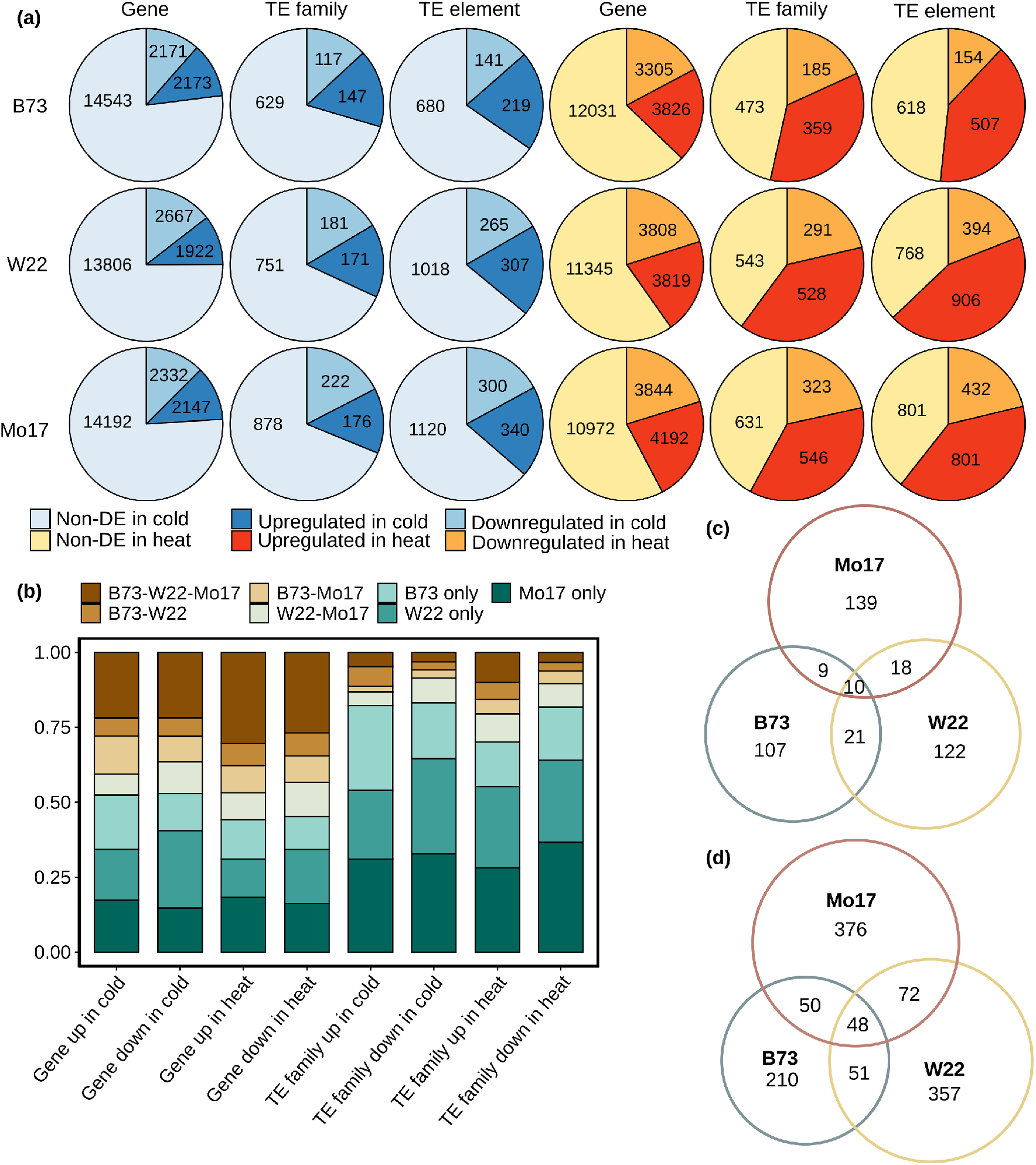
Expression changes for genes and TEs under cold and heat stress conditions. (a) The number of non-DE (but expressed), upregulated and downregulated genes, TE families and TE elements is shown for cold and heat conditions relative to control. (b) The consistency of up- or down-regulation was investigated for the non-redundant set of genes or TE families that are up- or down-regulated in at least one genotype. This analysis only included the set of genes and TE families that are present in all three genomes. The proportion of genes or TE families that are up-regulated in any combination of the three genotypes is shown. The overlap between shared TE families that are upregulated among three genotypes in (c) cold condition or in (d) heat condition is shown.

### Genotypic variation for TE responses to abiotic stress

We proceeded to evaluate how consistently the same TE families would respond to abiotic stress in multiple maize inbreds. Clustering of the differentially expressed (DE) genes or TE families (Figure S3) reveals differences in the consistency of response to abiotic stress for genes and TEs. While the genes are well clustered by treatment and reveal many examples of common responses in all genotypes, there is much less evidence for consistent responses of the same TE families in all three genotypes (Figure S3). While there are many genes that are consistently up- or down-regulated in multiple genotypes, the majority of TE families only show altered expression in one of the three genotypes (Figure 1b). A comparison of TE families that are present in all three genotypes reveals that only a subset of families were consistently up-regulated in all three genotypes (Figure 1c and 1d). There are 10 TE families with a consistent response to cold stress and 48 families with consistent response to heat stress. Within each genotype there are some TE families that are up-regulated in both cold- and heat-stress (Figure S4). There are 3 TE families (RLG08887, RLX10775 and RLX13504) that are up-regulated in all genotypes in both cold and heat stress conditions. These TE families may have a more global stress response.

### Trade-offs in per-family and per-element analyses of transposon expression

The analysis of per-family TE expression is quite useful to capture all transcripts arising from TEs, even if they are highly repetitive and map to many genomic locations. For the TE families that are up-regulated in B73 in heat or cold stress conditions, we assessed the proportion of uniquely mapping reads within the family (Figure 2a and 2b). Very similar results are observed for up-regulated TE families in W22 or Mo17 (Figure S5). This highlights the variability in the potential to capture expression using unique mapping reads compared to per-family estimates that use both unique and multi-mapping reads. In some TE families there are few or no unique mapping reads and thus a per-element expression analysis cannot identify individual members of these families. However, in other cases the majority of reads derived from a family can be uniquely mapped. While there is certainly value in characterizing all of the TE families that change in expression, the use of per-family expression estimates limits our ability to investigate the factors associated with expression of specific elements. Therefore, we performed a per-element expression analysis that only utilizes unique mapping reads to monitor expression of individual TEs. This approach often results in under-estimates of expression as multi-mapping reads are omitted. The analysis of per-element expression reveals similar proportions of up- and down-regulated elements when compared to per-family estimates (Figure 1a) and generates similar clustering of samples (Figure S1).

**Figure 2.**
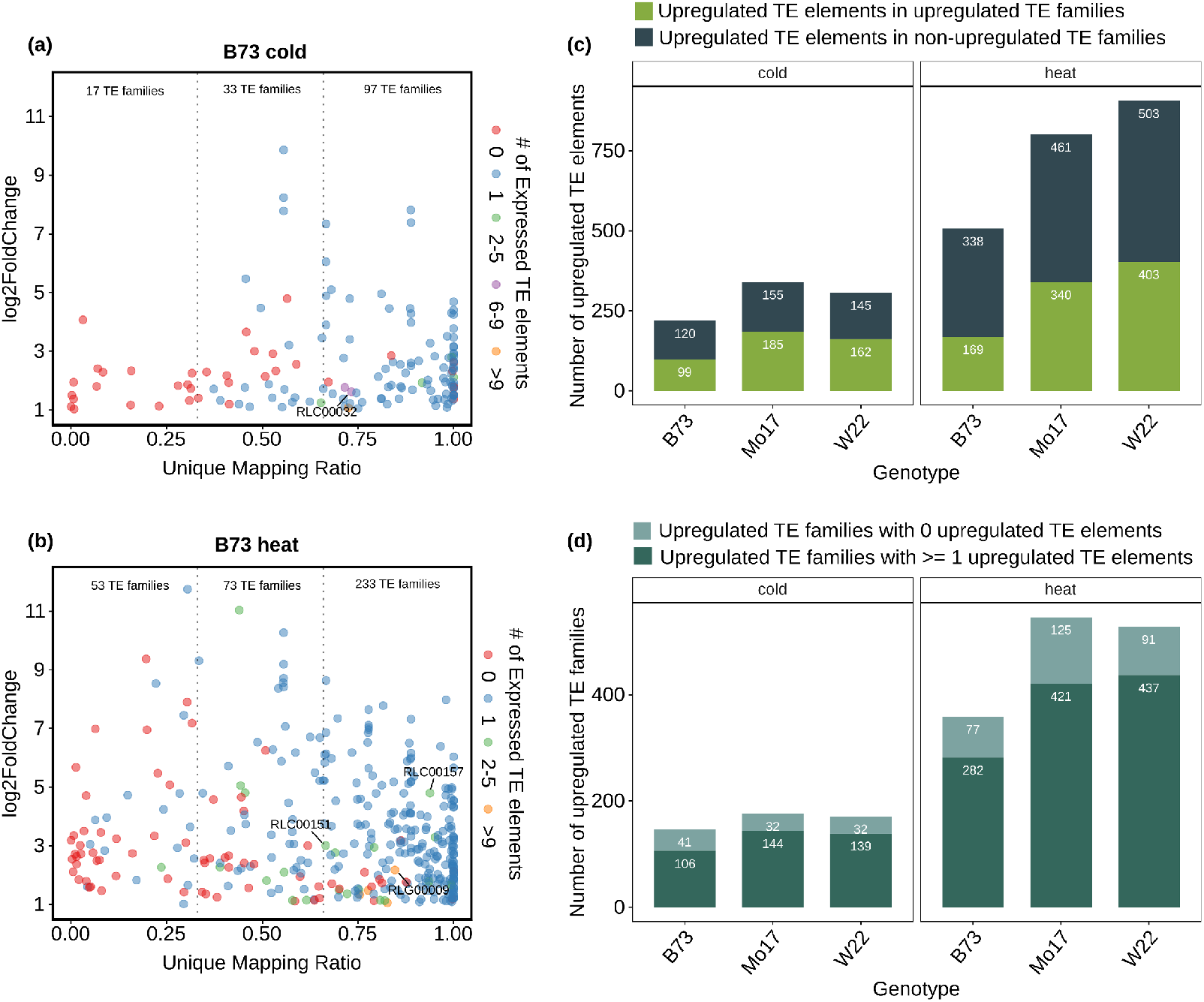
Comparison of per-family and per-element analyses of TE expression. (a) For all B73 TE families that are upregulated in cold condition the proportion of unique mapping reads attributed to the family was determined. The plot compares the expression change relative to control (y-axis) and proportion of unique mapping reads in cold stress (x-axis) for each upregulated TE family in B73. The color of each data point indicates the number of elements that are detected as expressed (> 1 CPM) in cold experiment. The number of TE families with <⅓, ⅓-⅔, or >⅔ unique mapping reads is shown. The upregulated TE families that are shared among three genotypes (B73, W22 and Mo17) with > ⅔ unique mapping ratio and >= 3 expressed TE elements were labeled. (b) The same information is presented for upregulated TE families under heat condition. (c) In each genotype the unique mapping reads were utilized to identify upregulated TE elements. These upregulated TE elements were compared to the results using multi-mapping reads to monitor per-family TE expression. The plot indicates the number of upregulated TE elements that in TE families that are significantly upregulated and the number of upregulated elements that are in TE families that were not identified as significantly up-regulated. (d) The number of significantly upregulated TE families with at least 1 or with 0 upregulated elements (based on unique mapping reads) is shown.

A comparison of the TE families and elements with altered expression reveals only partial overlap (Figure 2c and 2d). Only ~50% of the up-regulated TEs from the per-element analysis using unique mapping reads are within TE families that are classified as up-regulated (Figure 2c). This can occur for a variety of reasons. The most common reason is likely that the TE family is expressed in both control and stress conditions, but only some elements in the family are up-regulated and this does not generate enough difference to be classified as significant. We also assessed the proportion of up-regulated TE families that contain at least one up-regulated element (Figure 2d). While the majority of up-regulated TE families contain an up-regulated element there are 20-35% of the families that do not have any up-regulated elements. This is likely because some of these families are highly repetitive and we were not able to quantify individual elements within a family. For the rest of the analyses in this study, we largely focus on per-element expression estimates since this allows us to monitor factors associated with a specific genomic locus.

### Partial transcripts of TEs often arise near unmethylated TE regions

Our analyses that assess transcript levels for TE families or elements are based on counting reads that align to transposons. The presence of transcripts derived from TEs does not necessarily imply production of functional transcripts containing a full ORF that could provide transcriptional activity. However, visual inspection of the transcripts reveal that in many cases the transcripts are only observed for a part of the TE (Figure 3a-c). This suggests that many of the transcripts that occur following a stress treatment (or in control conditions) represent activity from internal cryptic promoters that may not result in any actual transposition events.

**Figure 3.**
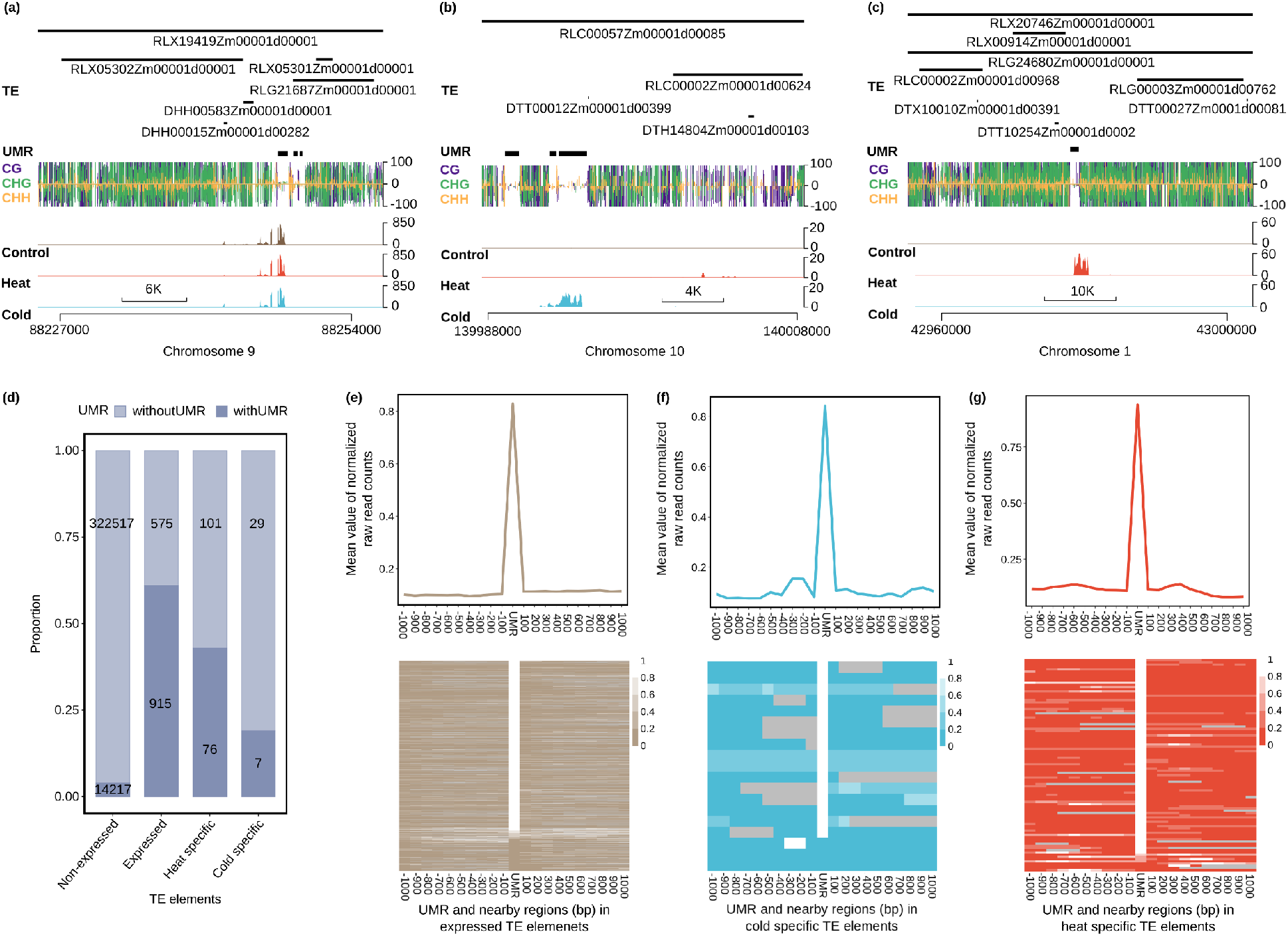
Localized expression within TEs often occurs near UMRs. (a-c) Examples of localized expression of a region within a TE are shown for three genomic regions. In (a) is an example of expression that occurs within/near a UMR in control and stress conditions. The examples in (b) and (c) show examples of cold-specific and heat-specific expression of TEs. In each plot the RNA-seq values are the average of all three biological replicates and the DNA methylation data is based on WGBS of the control sample and are visualized using the R package trackViewer (Ou & Zhu, 2019). (d) The proportion of TE elements with and without UMRs in non-expressed, expressed, heat-specific and cold-specific categories (see Methods for heat and cold-specific TE elements). Number in each class was indicated in the bar plot; (e-f) The relative level of expression throughout TEs was examined relative to the UMRs for all expressed (e), cold-specific (f) or heat-specific (g) elements that contain a UMR. The expression value for each UMR and its 20 surrounding extended 100 bins were normalized from 0 to 1. The mean value of each region were drawn on the upper panel and normalized values for every UMR and its extended bins were drawn as the heatmap below. Any extended bins overlapped with another UMR will be labeled as gray in the heatmap.

We assessed whether there were factors that associated with the expression of portions of specific TEs. While TEs are generally highly methylated there are some unmethylated regions (UMRs) that occur within TEs (Crisp *et al.*; Noshay *et al.*, 2020). WGBS data was available for B73 control, cold and heat stress and was used to call UMRs (Figure S6). In general, the overlap between control and heat or cold samples was very similar to the overlap between two replicates of control samples (Figure S6a). A comparison of the UMRs in control and heat or cold revealed >97% overlap and the vast majority (>95%) of UMRs found only in control or stress conditions were due to missing data in the other samples rather than actual changes in methylation state (Figure S6b). TEs that contain a UMR are much more likely to be detected as expressed compared to elements without a UMR (Figure 3d). Even for TEs that are only expressed in heat or cold conditions there is still an enrichment for a UMR in DNA methylation data from plants grown in non-stressed conditions. This suggests that TEs may often contain cryptic promoters or regulatory elements that only have the potential for activity when they are unmethylated. In some cases, the lack of methylation does not lead to constitutive expression but instead results in competence for expression in specific conditions. The pattern of transcript accumulation within a TE is correlated with the presence of the UMR (Figure 3e, 3f and 3g). The majority of transcripts occur within or adjacent to the UMR. These analyses highlight the potential for chromatin state to influence the potential expression of TEs.

### Consistent and variable responses to abiotic stress among members of a TE family

Transcriptional activation of a TE family in response to abiotic stress could reflect consistent up-regulation of all members of a family or specific changes to some members of the TE family. We focused on the subset of TE families that have >2/3 unique mapping reads TE elements and assessed the per-element expression levels in these families to monitor whether response to stress is family-wide or element-specific. In some TE families (such as DTH00434 and RLC00157) the increase in expression was driven by high levels of expression of a single element (Figure S7). In other TE families (such as RLC00151, RLG00292 and DTH11270) the increase in expression was attributed to multiple members of the family (Figure S7).

When only some of the members of a family are up-regulated, it could reflect potential read-through expression or proximity to nearby stress-responsive enhancers. In contrast, when there is consistent upregulation for multiple members of the same TE family, it may reflect the presence of cis-regulatory elements within the TE that provide responsiveness to the stress condition. We sought to determine if there are TE families with consistent response in all genotypes and multiple members of the family that respond. We required that the TE family was up-regulated in all three genotypes, that there are >2/3 unique mapping reads for the family and that at least two elements of the TE family have increased expression. This resulted in the identification of 1 TE family for cold stress (RLG00032) and 3 families for heat stress (RLC00151, RLC00157 and RLG0009). These families with consistent responses of multiple elements may reflect conserved cis-regulatory elements within the TE that promote stress-responsive expression.

We sought to examine some of the potential explanations for why some members of a TE family would respond to an abiotic stress while others did not. One potential explanation could be sequence divergence among family members. We selected two LTR TE families that had at least 10 elements with several that were upregulated in B73 and generated sequence phylogenies based on the 5’ LTR sequence (Figure S8). The elements that are up-regulated do not necessarily represent a specific subset of related sequences. This suggests that factors beyond sequence identity influence the responsiveness of individual elements.

An alternative possibility is that the potential to have transcriptional activation is a chromatin property rather than a sequence (genetic) property. We assessed chromatin variation in specific elements for the set of up-regulated TE families in B73 that have at least 2/3 of unique-mapping reads and >= 2 expressed TE elements. We assessed the proportion of elements with UMRs among the TEs in these families that are up-regulated compared to those that are not (Figure 4a and 4b). Within these families we find strong enrichments for UMRs within elements that are expressed in control or stress conditions compared to silent TEs. This suggests the potential for a TE to become transcriptionally active is dependent upon chromatin states that are present in control plants that are not expressing the TE.

**Figure 4.**
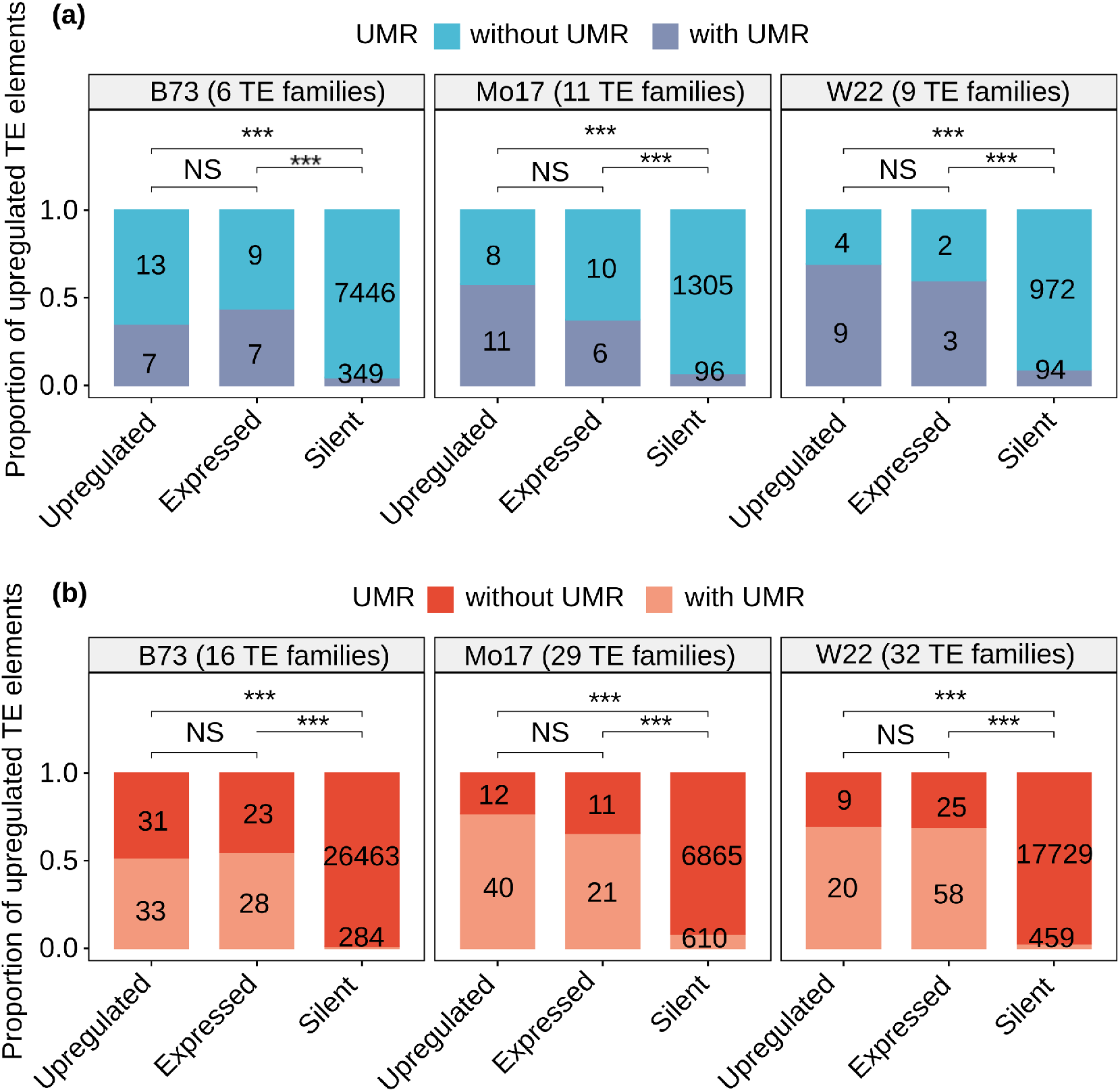
Methylation contributes to expression potential among members of TE families. We focused on a set of families (number of families indicated by genotype label) that exhibit up-regulation in cold stress that also have at least ⅔ of the expression derived from unique-mapping reads. The elements in these families were split into three groups; Expressed (>1 CPM but not significantly upregulated), Upregulated (log2foldchange > 1 and adjust p-value <= 0.05 for per-element expression), and Silent. For each of these three groups we assessed the number of elements within and without UMRs. Fisher exact test was performed to compare upregulated and one of other two categories (***: p-value < 0.001; NS: not significant). In (b) the same analysis was performed for the set of TE families with >⅔ unique mapping reads that are upregulated in heat stress.

### Documenting the basis for variable responses among genotypes

The observation that different genotypes exhibit quite distinct sets of TE families or TE elements with response to abiotic stress could reflect differences in the specific TEs present in each genotype or regulatory differences at shared elements. In order to address this in more detail, we focused on the per-element TE expression changes from TE families with significant up-regulation. We relied upon a prior set of TE polymorphism calls made using synteny blocks for these genomes (Anderson *et al.*, 2019a) and performed pairwise contrasts for the TEs that are up-regulated in each genome. Among the up-regulated B73 TEs with resolved presence/absence calls in the compared genome, the largest set of them are absent in the other genome (Figure 5a). Of the TEs that are present in both genomes a subset shows similar up-regulation in both genomes (purple in Figure 5a), but many of them are not changed (green in Figure 5a) or not expressed in the other genome (orange in Figure 5a). Generally similar proportions are seen in each of the comparisons of genomes (Figure 5b and 5c; Figure S9). This suggests that the differences in TE responses are driven both by polymorphic TE content and by regulatory variation of shared TEs.

**Figure 5.**
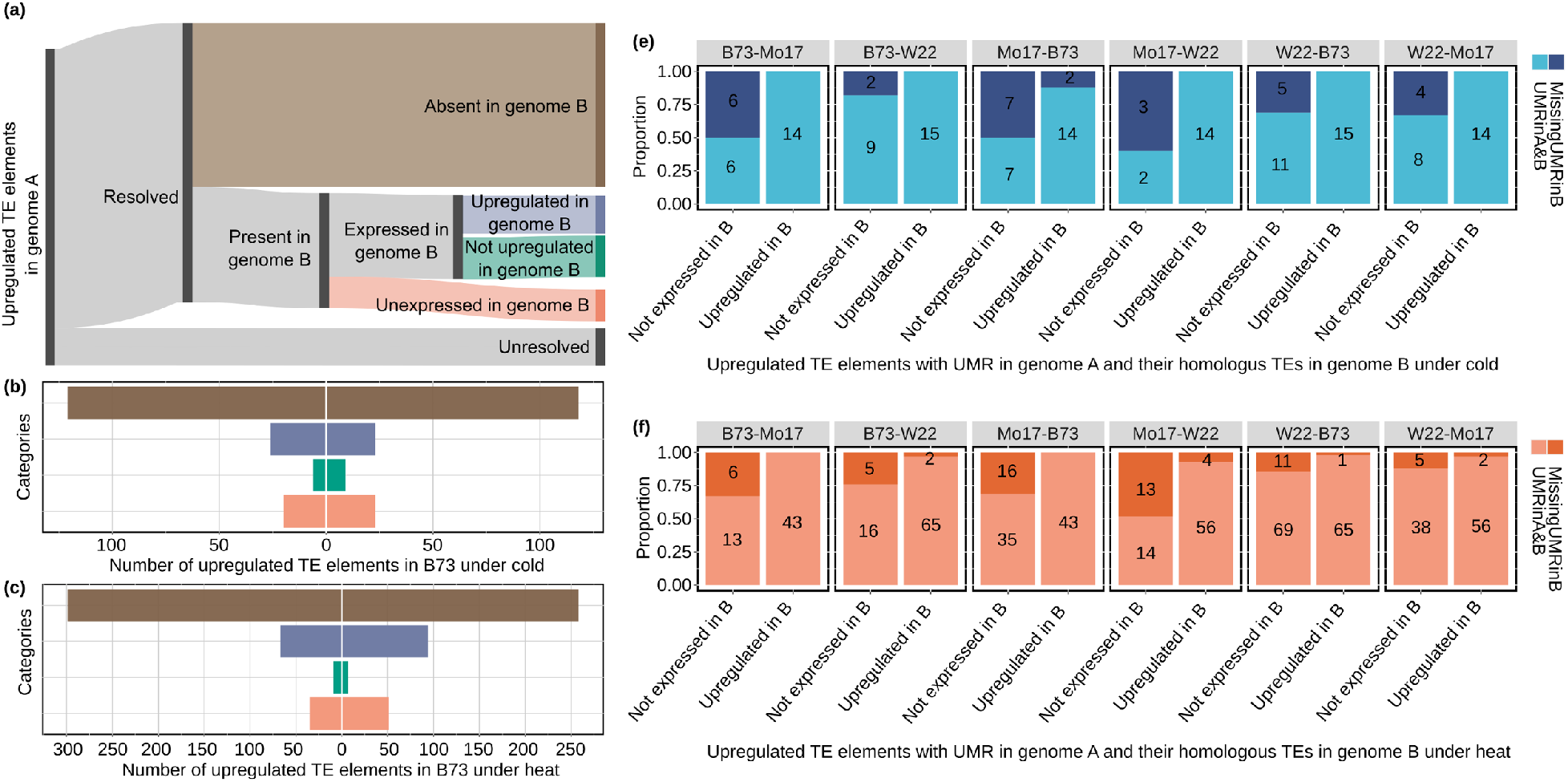
Assessing the factors that contribute to variable responses between genotypes. (a) A schematic diagram showing how upregulated TE elements in genome A are classified into different categories in a comparison with genome B. The presence/absence (or unresolved) nature of TE elements in one genome relative to another is from Anderson et al. 2019. For the subset of TE elements present in both genomes the expression in genome B is classified as silent (orange), significantly upregulated (blue), expressed but significantly upregulated (green). The number of TE elements in categories defined in panel a under cold condition when B73 was served as genome A and compared to W22 or Mo17 as genome B. In (c) a similar plot is shown for comparisons of B73 heat upregulated elements with Mo17 or W22. (d-e) The upregulated TE elements in genome A with a UMR were used to assess whether loss of UMRs might contribute to loss of stress-responsive expression. For each contrast of two genomes we investigated the proportion of elements with or without UMRs in genotype B for shared elements that are upregulated in both genomes compared to elements that are up-regulated in one genome but not expressed in the other.

We further examined the shared TEs that are up-regulated in one genome but not expressed in the other. We hypothesized that these could reflect examples of chromatin variation in which there are unmethylated regions in the genotype that can be activated but fully methylated chromatin in other genotypes which limits the potential for expression in that genotype. In each genotype (genotype A), we identified the subset of TEs that are (1) up-regulated in response to heat or cold, (2) contain a UMR and (3) have a shared insertion in the genotype being compared (genotype B). We then classified whether the TE was also up-regulated in genotype B or was silent in genotype B. We then evaluated the proportion of these cases in which the UMR was present in genotype B compared to cases in which the TE is fully methylated in genotype B. In all pairwise contrasts we found that when the TE was up-regulated in response to heat or cold stress, the UMR was usually (>90% of examples in each case) found in both genotypes (Figure 5c and 5d). In contrast, for examples in which the TE was not expressed in genotype B there was a much higher rate of UMR loss (Figure 5c and 5d). This suggests that one reason for the polymorphic responsiveness of shared TE elements in the different genotypes is due to differences in chromatin that influence the potential for transcriptional activation.

## Discussion

TEs are ubiquitous components of many eukaryotic genomes but we are still developing an understanding of how they contribute to the transcriptome. In this study we have focused on a replicated analysis of the transcriptome of three maize inbreds in three environmental conditions. Our observations highlight the potential roles of both genetic and epigenetic variation in contributing to expression variation of different transposable elements.

### Trade-offs in per-family and per-element analyses of TE expression

The highly repetitive nature of many TE families results in complications for the analysis of TE expression as well as the analysis of chromatin state. In addition, the lack of precise annotation of TE transcripts can also complicate the analysis of TE expression. In this study we have employed two complementary approaches to monitor TE expression. One approach uses both unique and multi-mapping reads to monitor the expression of TE families. This casts a broad net that can identify the majority of transcripts arising from TEs, even in highly repetitive families. However, this type of analysis does not identify the specific elements, or genomic locations, that contribute to expression. This limits our ability to assess whether many members of the same family exhibit a coordinate response and also limits our ability to assess chromatin state at expressed elements. It is worth noting that the issue of multi-mapping also complicates comparisons of chromatin state. Many TEs cannot be assessed for methylation or the presence of UMRs as they are too repetitive. This complicates some of our analyses of the presence of UMRs in different categories of TEs as the lack of a UMR may reflect the true absence of a UMR or it could miss data due to lack of unique alignments.

### Genomic variation contributes to polymorphic responses

A comparison of genic and TE transcriptome responses to heat/cold stress revealed that there is more variability among genotypes for TE responses than for gene expression responses. A detailed examination of the TEs that respond to heat or cold stress in one genotype revealed that many of these specific TEs are not present in the same genomic location in the other lines. In each contrast over half of the TEs that are up-regulated in one genome lack a syntenic insertion in other genomes. This suggests that much of the variable transcriptional response in a comparison of genomes actually is a result of genome content differences. This observation highlights a challenge of comparing transcriptional responses of TEs. These studies are highly reliant upon the genome being used for alignment. Many studies that compare transcriptional responses rely upon alignments to a common reference genome. However, the highly variable nature of TE insertions among maize lines will result in difficulties in comparing the expression responses.

### Chromatin state influences TE expression potential

While variation in genome content explains some of the variation for TE expression responses there are also many shared insertions in two genomes that exhibit distinct expression responses. While there certainly can be examples of trans-regulatory differences that could affect abiotic stress response, there are more examples of cis-regulatory variation than trans-regulatory variation (Waters *et al.*, 2017). However, shared TE insertions in two genomes are expected to have largely similar sequences. We hypothesized that differences in chromatin state may explain some of the variation in transcriptome responses among genotypes. We found that TEs that are expressed are highly enriched for detectable UMRs in maize. DNA methylation remains virtually identical in the control and stress conditions with very few novel UMRs in cold or heat stressed samples. However, many of the TEs that exhibit expression only in heat or cold stress contain UMRs in control conditions. This suggests that the lack of DNA methylation allows for potential regulatory elements to respond to the presence of trans-acting factors that induce heat- or cold-responsive expression.

A comparison of TEs that show variable response in two genomes finds many examples in which there is evidence for changes in chromatin state. Often, the failure to respond to an environmental stress in a second genotype is accompanied by the loss of the UMR. This region exhibits variable methylation between genotypes and the highly methylated genotype losses the potential for expression.

The role of chromatin state could also play an important role in explaining the variability of responses of different TEs within a family. With a TE family many of the elements can have highly similar sequences. While there are a small set of families that show coordinate expression changes of many elements in the same family, the majority of TEs that are activated by heat or cold stress reflect a small number of elements within the family. When we assessed the presence of UMRs in these elements we found that the elements that are expressed or up-regulated in stress conditions are much more likely to contain UMRs. This suggests that the presence of a UMR is likely important for providing expression potential for TEs.

## Acknowledgements

We are grateful to Peter Hermanson for generating the necessary plant materials and samples for these experiments. The Minnesota Supercomputing Institute (MSI) at the University of Minnesota provided computational resources that contributed to this research. Research on this project was supported by grants from the NSF Plant Genome Research Program (IOS-1934384), USDA-NIFA (2016-67013-24747) and by Hatch funding front he Minnesota Agricultural Experiment Station (MIN 71-068). PAC is the recipient of an Australian Research Council Discovery Early Career Award (project number DE200101748).

## Author contributions

Z.L., S.N.A. and N.M.S. conceived the experiments. S.N.A. and T.A.E. grew plants and collected RNA samples. Z.L., J.M.N. and P.A.C. analyzed data. Z.L., S.N.A., J.M.N., P.A.C. and N.M.S. drafted the manuscript. All authors discussed the results and contributed to the final manuscript.

## Competing interests

The authors declare that they have no competing interests.

## Supporting information

**Figure S1. Principal component analysis for all replicates under control, cold and heat conditions for B73, W22 and Mo17 using expressed genes, TE families or TE elements.** For genes, TE families or TE elements expression was defined as CPM > 1 in at least 5 libraries. Only the first 2 principal components were used for visualizations. The percent of explained variation for each component was labeled for PC1 and PC2.

**Figure S2. The distribution of TE families sizes is shown for different groups of TEs.** TE family size was classified as 1 (red), 2-9 (tan), 10-50 elements (light blue) and >50 elements (dark blue). The number of families with each size is shown for up-regulated TE families for each genotype and condition. In addition, the number of families of each size is shown genome-wide (total) or for expressed families (Exp).

**Figure S3. Transcript abundance clustering of expressed genes and TE families.** (a) Heatmap for gene expression abundance. Only genes with mean CPM > 1 were shown and each gene per row was normalized for visualization; (b) A heatmap of clustered per-family TE expression (including both unique and multiple-mapping reads) was generated for expressed TE families. Only TE families with mean CPM > 1 were shown and each row was normalized for visualization.

**Figure S4. Overlap of up-regulated TE families in heat and cold conditions for each of the three genotypes.** (a) B73; (b) W22; (c) Mo17.

**Figure S5. Comparison of per-family and per-element analyses of TE expression for W22 (a-b) and Mo17 (c-d).** For all TE families that are up-regulated in cold condition the proportion of unique mapping reads attributed to the family was determined. The plot compares the expression change relative to control (y-axis) and proportion of unique mapping reads in cold stress (x-axis) for each up-regulated TE family. The color of each data point indicates the number of elements that are detected as expressed (>1 CPM) in cold condition. The number of TE families with <⅓, ⅓-⅔, or >⅔ unique mapping reads is shown. The W22 up-regulated families were assessed in both cold (a) and heat (b) condition. Similarly, both cold (c) and heat (d) condition families are shown for Mo17.

**Figure S6. Comparisons of unmethylated regions.** (a) Overlap of unmethylated regions (UMRs) in two independent maize seedling leaf control samples of B73, "Rep 1" and "Rep 2", cold and heat treated leaves. The percent of UMRs uniquely identified in one of the samples is listed in parantheses and venn diagrams are not in proportion; (b) For UMRs uniquely identified in one of the samples in a, the methylation data (methylation domain type) in the corresponding sample is displayed. The majority of "unique" UMRs identified in stress conditions occur because of missing data one of the samples.

**Figure S7. Monitoring the contribution of individual elements to per-family increase in expression.** Six example TE families with >⅔ unique mapping ratio were selected to monitor the relative contribution of individual elements to total expression. Each individual subplot indicates a TE family and shows relative expression of elements in control, cold and heat conditions. The same color within each subplot indicates the same TE element. The mean CPM value of all replicates under each condition was represented on the y-axis. (a) The cold upregulated TE family RLG00037; (b) The cold upregulated TE familly DTH000434; (c) The heat upregulated TE family RLC00157; (d) The heat upregulated TE family RLC00151; (e) The TE family RLG00292 that is upregulated in both cold and heat; (f) The TE family DTH11270 that is upregulated under both cold and heat.

**Figure S8. Phylogenetic trees of TE elements in four selected upregulated LTR TE families under heat stress in B73.** (a) The 5’ LTR sequence per TE element within RLC00157 TE family was used to construct the phylogenetic tree. The upregulated, expressed and UMR containing status was labeled on the right side per TE element; (b) RLC00151 TE family. Two solo LTRs within this TE family were not considered.

**Figure S9. Number of TE elements in categories defined in Figure 5a.** (a) W22 was served as genome A and compared to W22 or Mo17 as genome B under cold condition; (b) W22 was served as genome A and compared to W22 or Mo17 as genome B under heat condition; (c) Mo17 was served as genome A and compared to W22 or B73 as genome B under cold condition; (d) Mo17 was served as genome A and compared to W22 or B73 as genome B under heat condition.

---

**Table S1.** GO enrichment for upregulated genes in three maize genotypes under cold and heat stress.

**Table S2.** Upregulated TE families under cold and stress conditions in three maize genotypes.

## Notes

### Competing Interest Statement

The authors have declared no competing interest.

## References

Anderson SN, Stitzer MC, Brohammer AB, Zhou P, Noshay JM, O’Connor CH, Hirsch CD, Ross-Ibarra J, Hirsch CN, Springer NM. 2019a. Transposable elements contribute to dynamic genome content in maize. The Plant journal: for cell and molecular biology 100: 1052–1065.

Anderson SN, Stitzer MC, Zhou P, Ross-Ibarra J, Hirsch CD, Springer NM. 2019b. Dynamic patterns of transcript abundance of transposable element families in maize. G3: Genes, Genomes, Genetics 9: 3673–3682.

Anders S, Pyl PT, Huber W. 2015. HTSeq--a Python framework to work with high-throughput sequencing data. Bioinformatics 31: 166–169.

Benoit M, Drost H, Catoni M, Gouil Q, Lopez-Gomollon S, Baulcombe D, Paszkowski J. 2019. Environmental and epigenetic regulation of Rider retrotransposons in tomato. PLoS genetics 15: e1008370.

Cavrak VV, Lettner N, Jamge S, Kosarewicz A, Bayer LM, Scheid OM. 2014. How a retrotransposon exploits the plant’s heat stress response for its activation. PLoS genetics 10: e1004115.

Crisp PA, Marand AP, Noshay JM, Zhou P, Lu Z, Schmitz RJ, Springer NM. 2020. Stable unmethylated DNA demarcates expressed genes and their cis-regulatory space in plant genomes. bioRxiv 2020.05.23.109744.

Emms DM, Kelly S. 2015. OrthoFinder: solving fundamental biases in whole genome comparisons dramatically improves orthogroup inference accuracy. Genome biology 16: 157.

Galindo-González L, Mhiri C, Deyholos MK, Grandbastien M-A. 2017. LTR-retrotransposons in plants: Engines of evolution. Gene 626: 14–25.

Grandbastien MA, Audeon C, Bonnivard E, Casacuberta JM, Chalhoub B, Costa AP, Le QH, Melayah D, Petit M, Poncet C, et al. 2005. Stress activation and genomic impact of Tnt1 retrotransposons in Solanaceae. Cytogenetic and genome research 110: 229–241.

Hirochika H. 1993. Activation of tobacco retrotransposons during tissue culture. The EMBO journal 12: 2521–2528.

Ito H, Gaubert H, Bucher E, Mirouze M, Vaillant I, Paszkowski J. 2011. An siRNA pathway prevents transgenerational retrotransposition in plants subjected to stress. Nature 472: 115–119.

Jiang N, Bao Z, Zhang X, Eddy SR, Wessler SR. 2004. Pack-MULE transposable elements mediate gene evolution in plants. Nature 431: 569–573.

Jiao Y, Peluso P, Shi J, Liang T, Stitzer MC, Wang B, Campbell MS, Stein JC, Wei X, Chin C-S, et al. 2017. Improved maize reference genome with single-molecule technologies. Nature 546: 524–527.

Katoh K, Standley DM. 2013. MAFFT multiple sequence alignment software version 7: improvements in performance and usability. Molecular biology and evolution 30: 772–780.

Kim D, Langmead B, Salzberg SL. 2015. HISAT: a fast spliced aligner with low memory requirements. Nature methods 12: 357–360.

Klopfenstein DV, Zhang L, Pedersen BS, Ramírez F, Vesztrocy AW, Naldi A, Mungall CJ, Yunes JM, Botvinnik O, Weigel M, et al. 2018. GOATOOLS: A Python library for Gene Ontology analyses. Scientific reports 8: 10872.

Krueger F. 2012. Trim Galore: a wrapper tool around Cutadapt and FastQC to consistently apply quality and adapter trimming to FastQ files, with some extra functionality for MspI-digested RRBS-type (Reduced Representation Bisufite-Seq) libraries. URL http://www.bioinformatics.babraham.ac.uk/projects/trim_galore/. (Date of access: 28/04/2016).

Kurtz S, Phillippy A, Delcher AL, Smoot M, Shumway M, Antonescu C, Salzberg SL. 2004. Versatile and open software for comparing large genomes. Genome biology 5: R12.

Lanciano S, Mirouze M. 2018. Transposable elements: all mobile, all different, some stress responsive, some adaptive? Current opinion in genetics & development 49: 106–114.

Li H, Handsaker B, Wysoker A, Fennell T, Ruan J, Homer N, Marth G, Abecasis G, Durbin R. 2009. The sequence alignment/map format and SAMtools. Bioinformatics 25: 2078–2079.

Lyons E, Pedersen B, Kane J, Freeling M. 2008. The Value of Nonmodel Genomes and an Example Using SynMap Within CoGe to Dissect the Hexaploidy that Predates the Rosids. Tropical Plant Biology 1: 181–190.

Makarevitch I, Waters AJ, West PT, Stitzer M, Hirsch CN, Ross-Ibarra J, Springer NM. 2015. Transposable elements contribute to activation of maize genes in response to abiotic stress. PLoS genetics 11: e1004915.

Negi P, Rai AN, Suprasanna P. 2016. Moving through the Stressed Genome: Emerging Regulatory Roles for Transposons in Plant Stress Response. Frontiers in plant science 7: 1448.

Noshay JM, Marand AP, Anderson SN, Zhou P, Guerra MKM, Lu Z, O’Connor C, Crisp PA, Hirsch CN, Schmitz RJ, et al. 2020. Cis-regulatory elements within TEs can influence expression of nearby maize genes. bioRxiv: 2020.05.20.107169.

Ou J, Zhu L. 2019. trackViewer: a Bioconductor package for interactive and integrative visualization of multi-omics data. Nature methods 16: 453–454.

Peschke VM, Phillips RL, Gengenbach BG. 1987. Discovery of transposable element activity among progeny of tissue culture–derived maize plants. Science 238: 804–807.

Price MN, Dehal PS, Arkin AP. 2010. FastTree 2--approximately maximum-likelihood trees for large alignments. PloS one 5: e9490.

Rabinowicz PD, Schutz K, Dedhia N, Yordan C, Parnell LD, Stein L, McCombie WR, Martienssen RA. 1999. Differential methylation of genes and retrotransposons facilitates shotgun sequencing of the maize genome. Nature genetics 23: 305–308.

Springer NM, Anderson SN, Andorf CM, Ahern KR, Bai F, Barad O, Barbazuk WB, Bass HW, Baruch K, Ben-Zvi G, et al. 2018. The maize W22 genome provides a foundation for functional genomics and transposon biology. Nature genetics 50: 1282–1288.

Stitzer MC, Anderson SN, Springer NM, Ross-Ibarra J. 2019. The Genomic Ecosystem of Transposable Elements in Maize. bioRxiv 2019.02.28.559922.

Sun S, Zhou Y, Chen J, Shi J, Zhao H, Zhao H, Song W, Zhang M, Cui Y, Dong X, et al. 2018. Extensive intraspecific gene order and gene structural variations between Mo17 and other maize genomes. Nature genetics 50: 1289–1295.

Vicient CM. 2010. Transcriptional activity of transposable elements in maize. BMC genomics 11: 601.

Waters AJ, Makarevitch I, Noshay J, Burghardt LT, Hirsch CN, Hirsch CD, Springer NM. 2017. Natural variation for gene expression responses to abiotic stress in maize. The Plant journal: for cell and molecular biology 89: 706–717.

Wessler SR. 1996. Turned on by stress. Plant retrotransposons. Current biology: CB 6: 959–961.

Wicker T, Sabot F, Hua-Van A, Bennetzen JL, Capy P, Chalhoub B, Flavell A, Leroy P, Morgante M, Panaud O, et al. 2007. A unified classification system for eukaryotic transposable elements. Nature reviews.Genetics 8: 973–982.

Wimalanathan K, Friedberg I, Andorf CM, Lawrence-Dill CJ. 2018. Maize GO Annotation-Methods, Evaluation, and Review (maize-GAMER). Plant direct 2: e00052.

Xi Y, Li W. 2009. BSMAP: whole genome bisulfite sequence MAPping program. BMC bioinformatics 10: 232.

Yu G, Smith DK, Zhu H, Guan Y, Lam TT. 2017. ggtree: an r package for visualization and annotation of phylogenetic trees with their covariates and other associated data. Methods in Ecology and Evolution 8: 28–36.

