## Supplementary figures and images for "Genetic and epigenetic contributions to variation in transposable element expression responses to abiotic stress in maize"

### Figure_S1

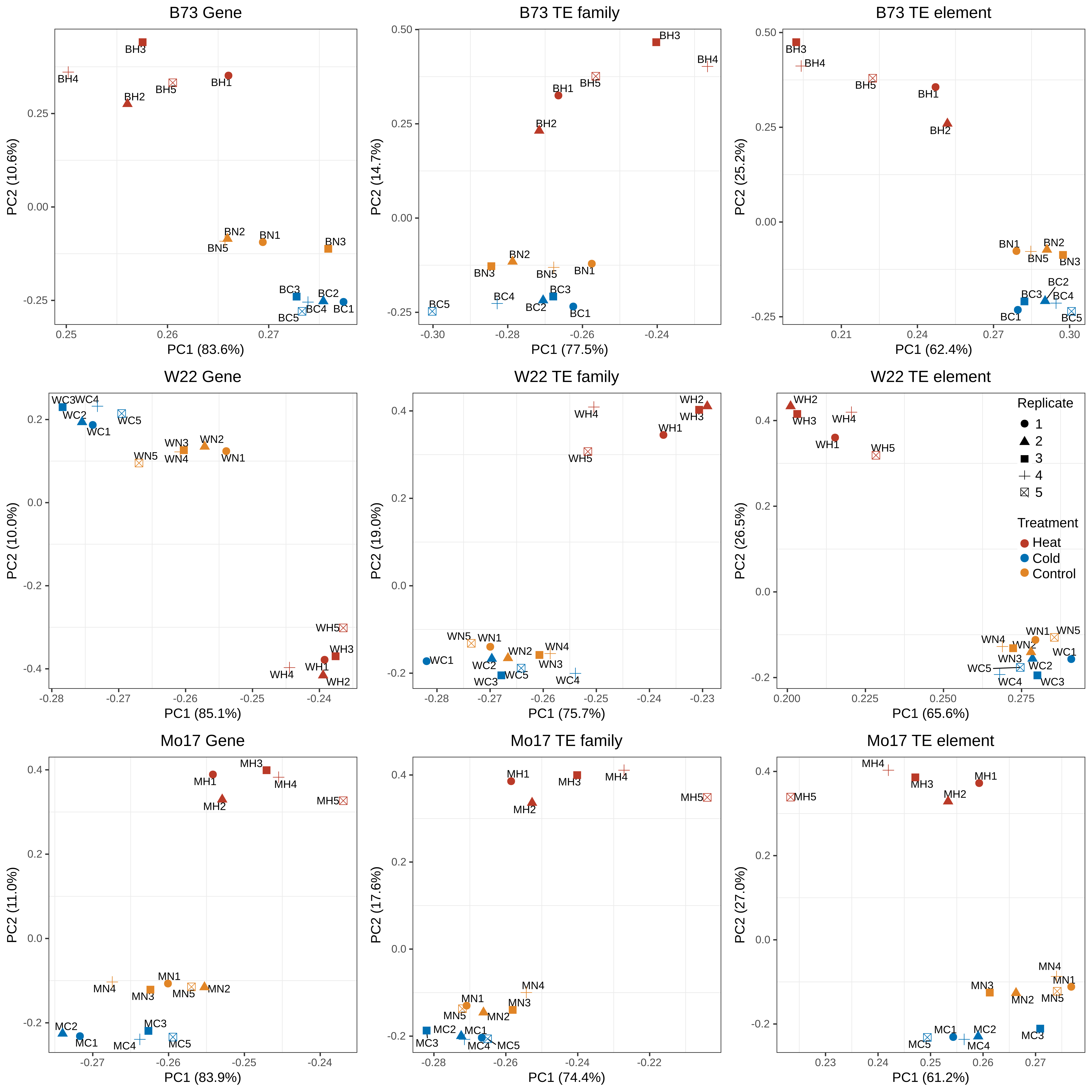

### Figure_S2

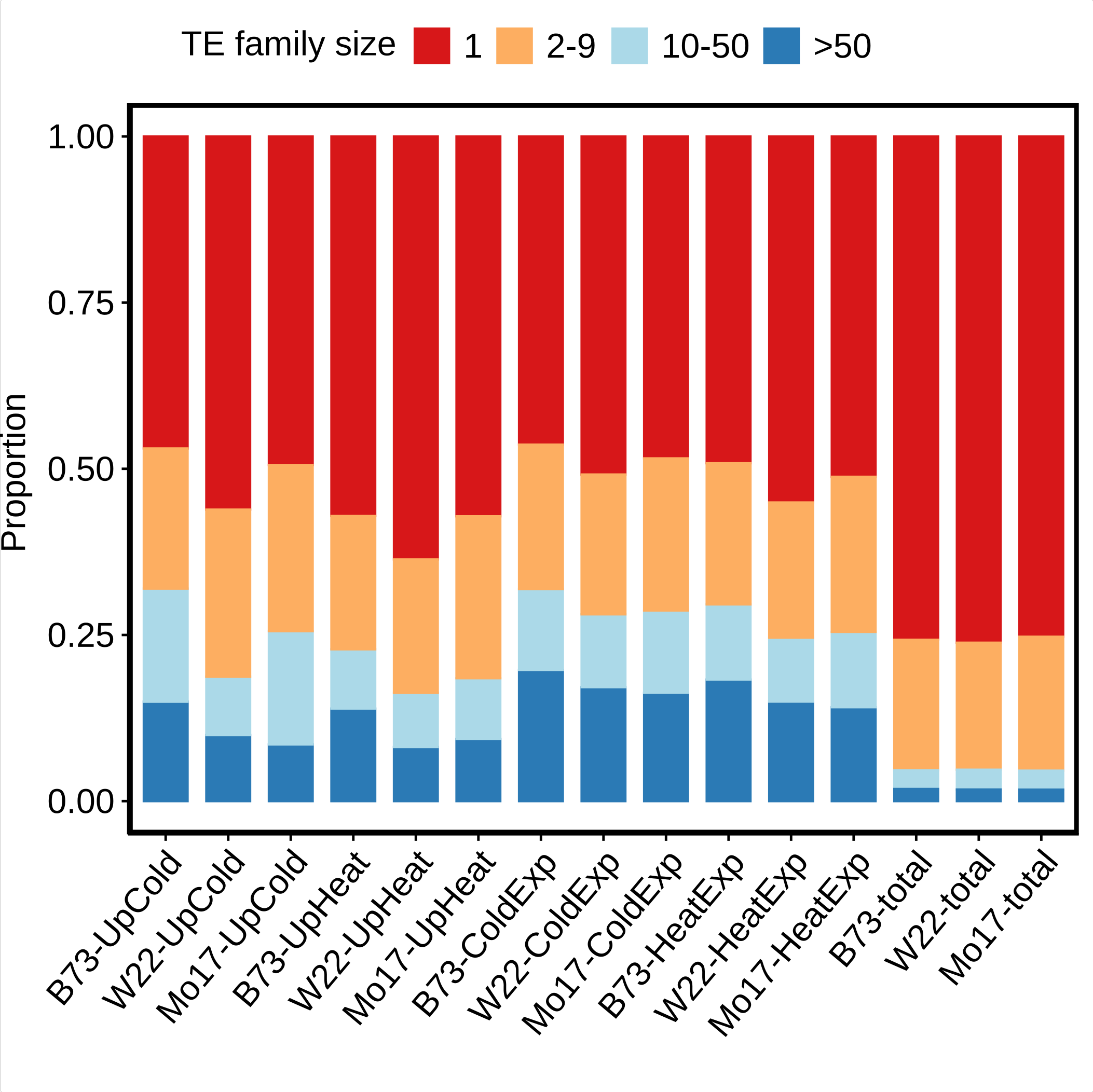

### Figure_S3

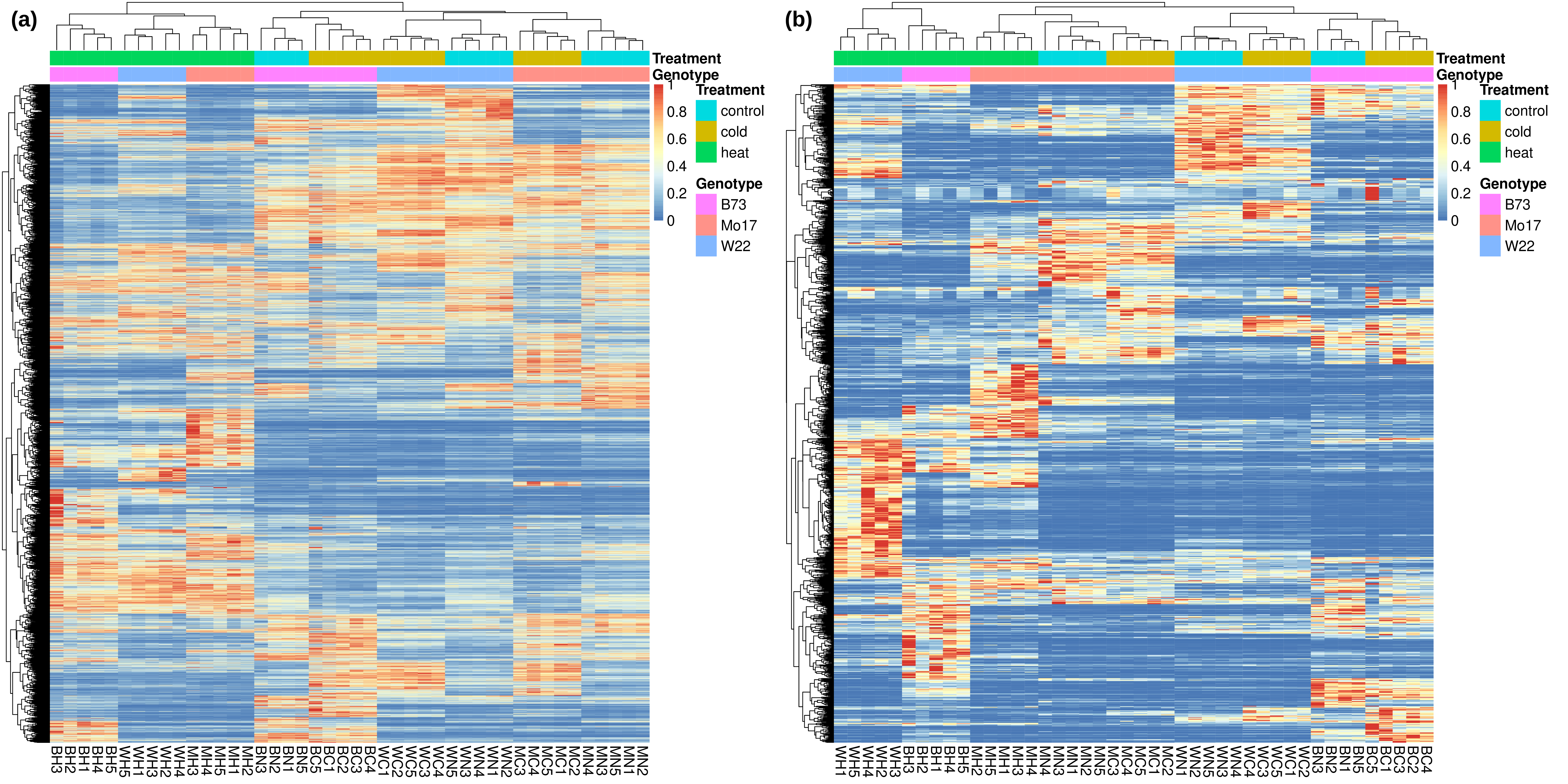

### Figure_S4

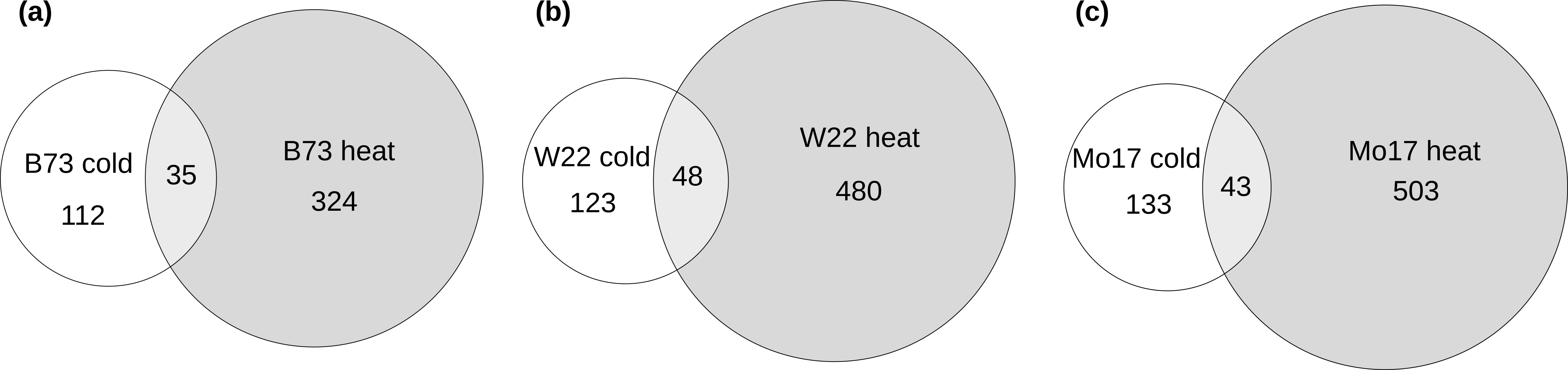

### Figure_S5

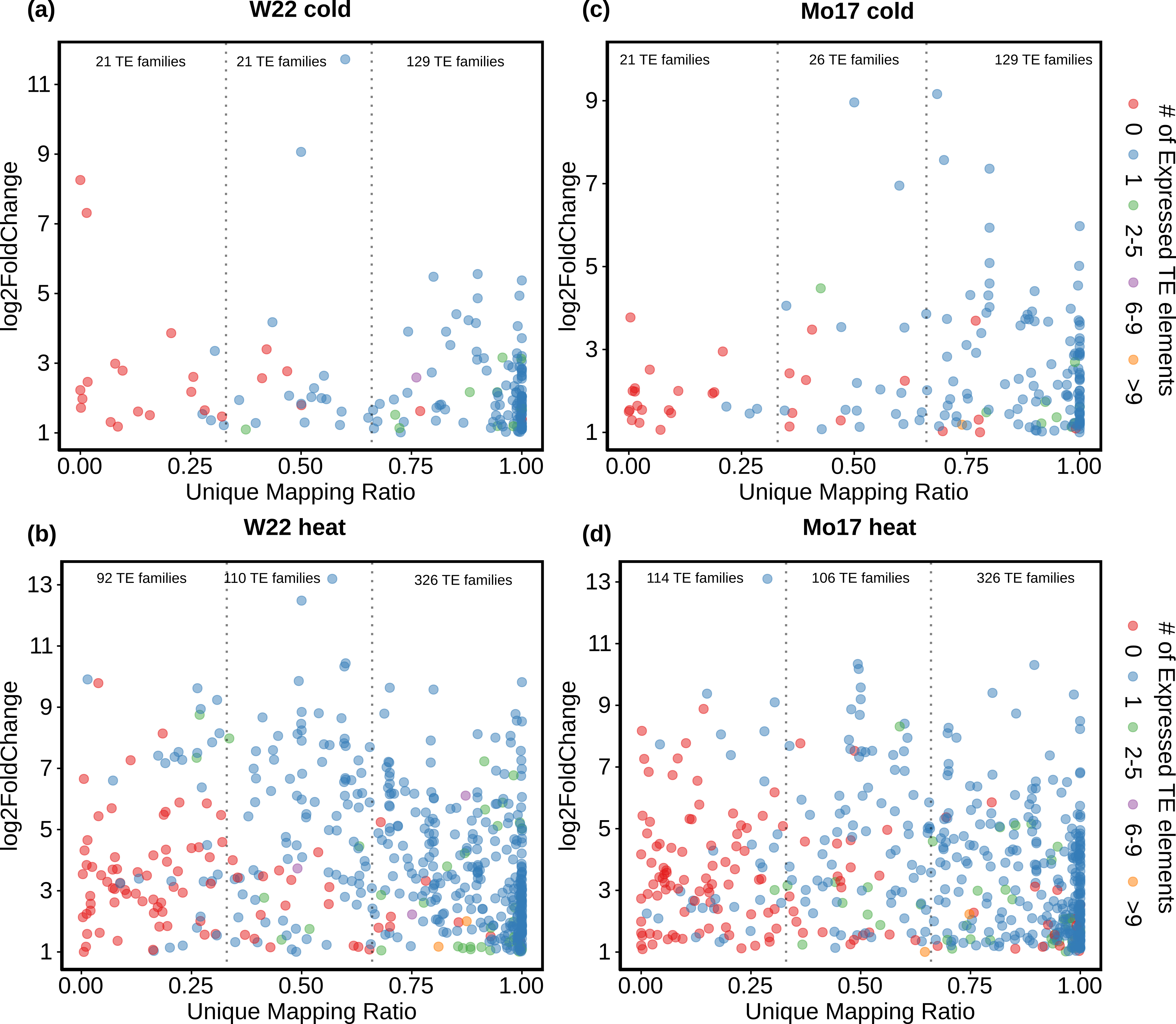

### Figure_S6

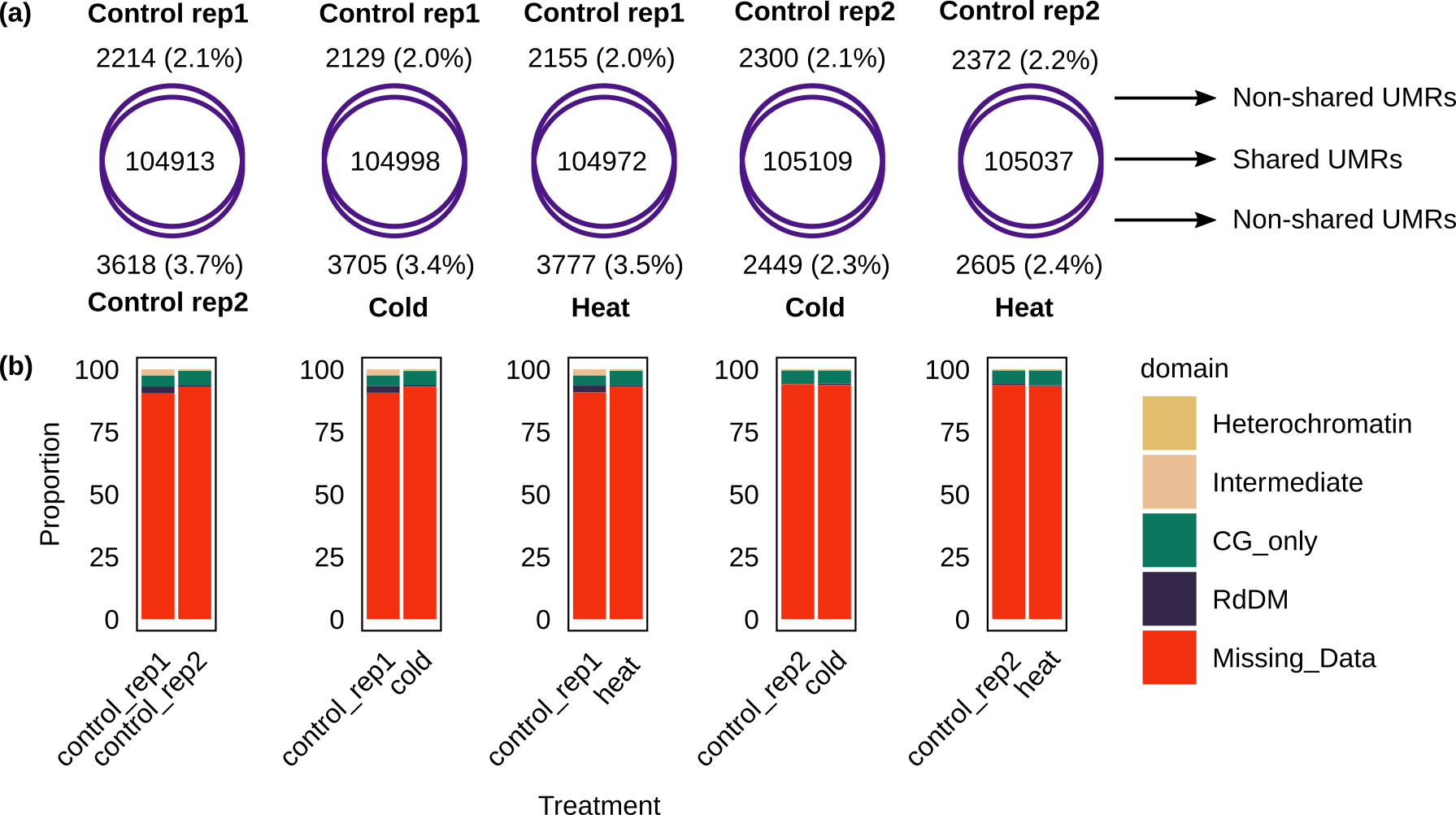

### Figure_S7

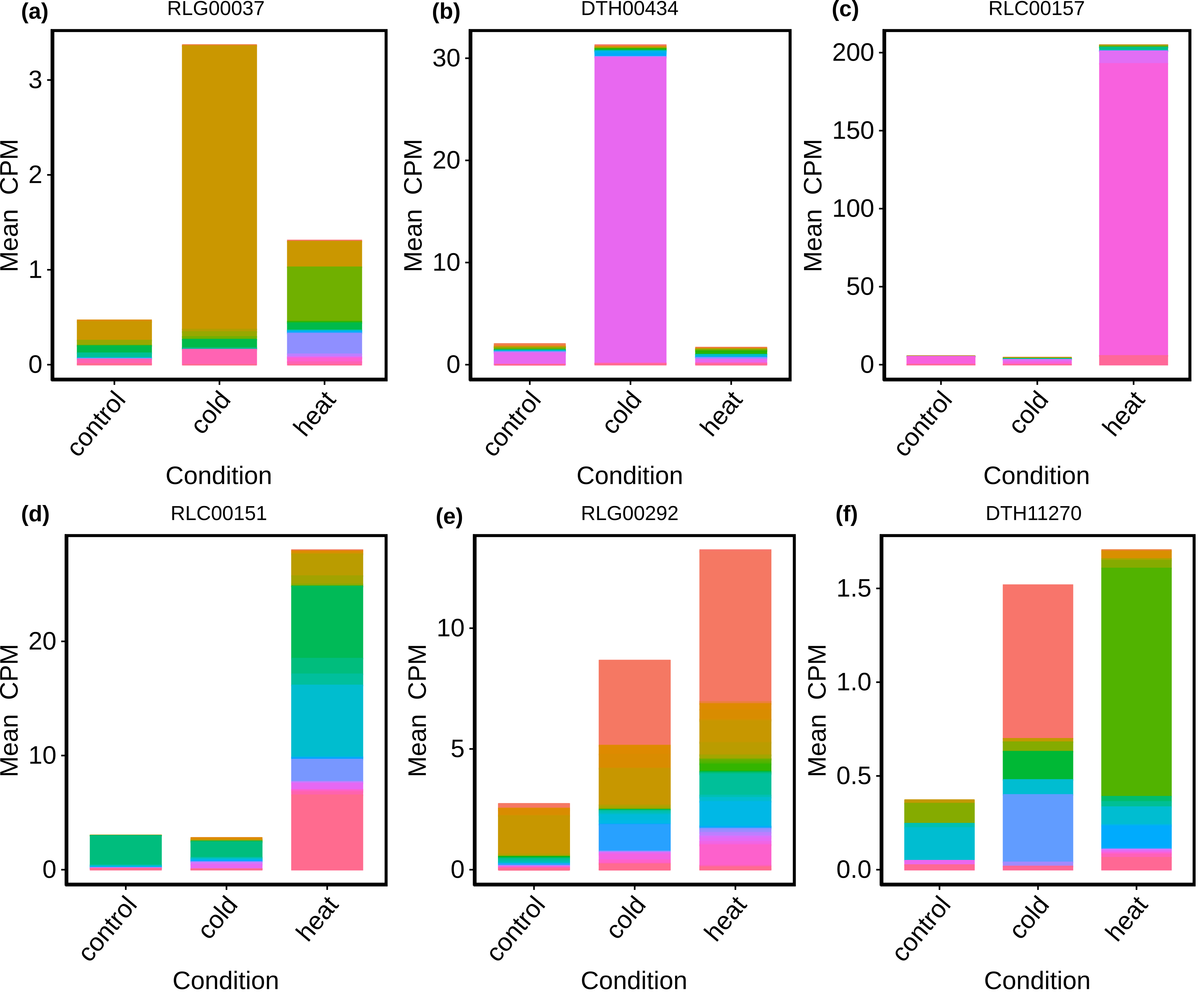

### Figure_S8

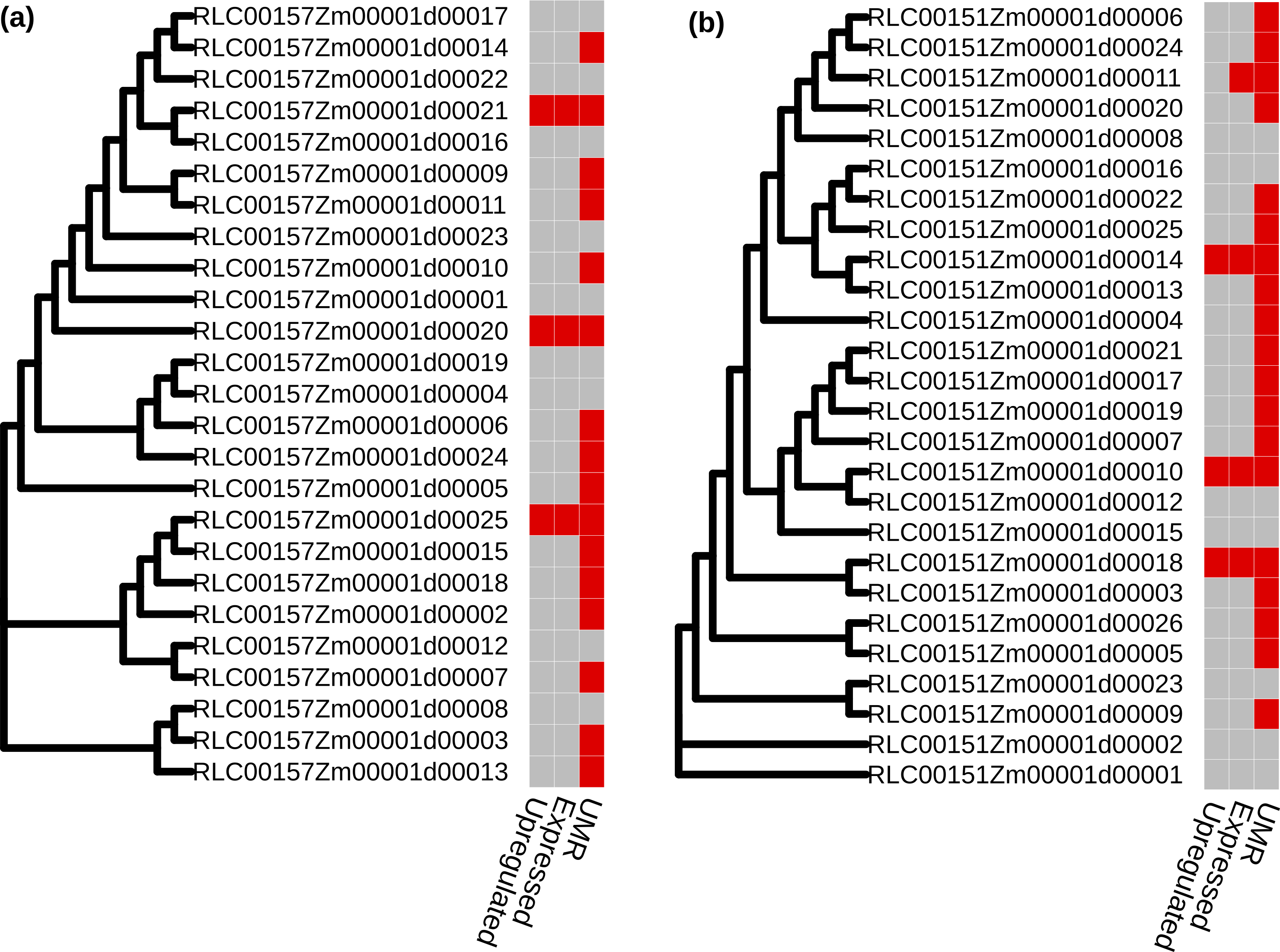

### Figure_S9

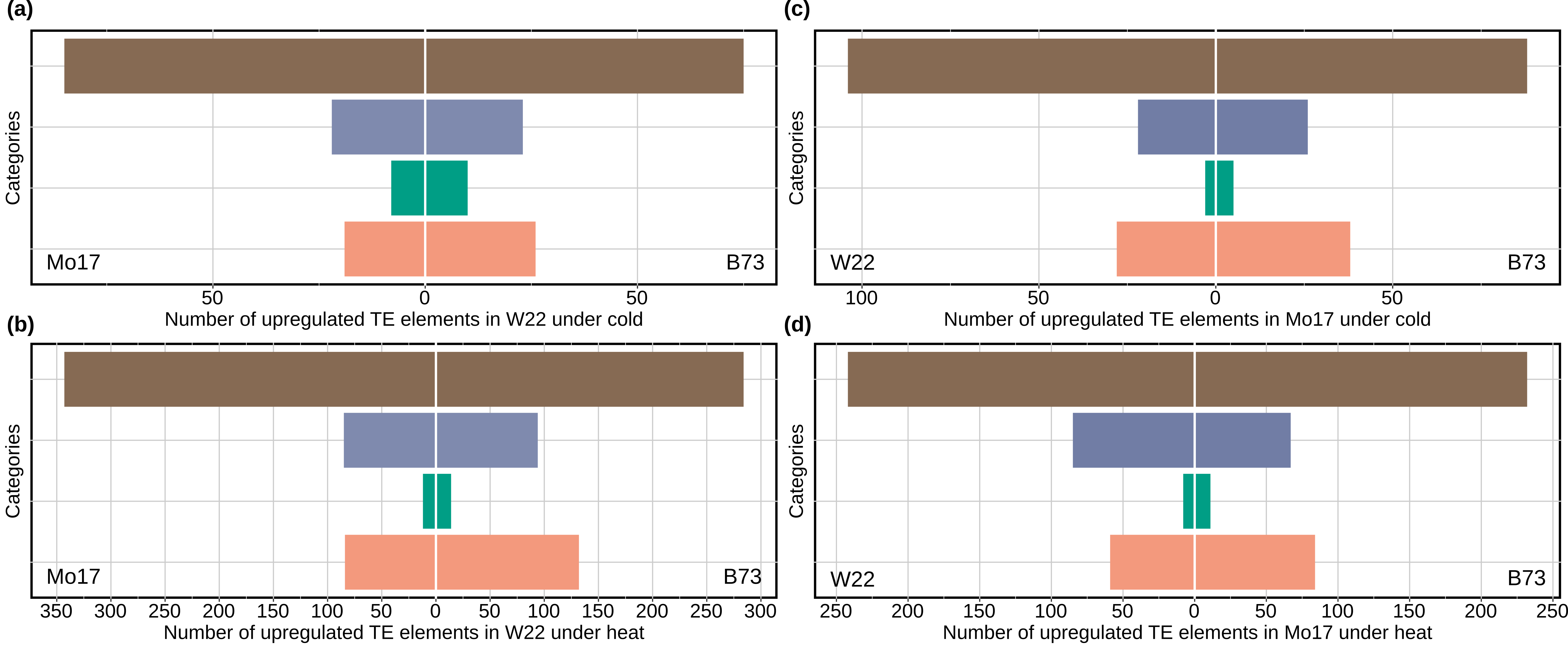
